# Parental histone distribution at nascent strands controls homologous recombination during DNA damage tolerance

**DOI:** 10.1101/2022.04.05.487148

**Authors:** Cristina González-Garrido, Félix Prado

**Affiliations:** Centro Andaluz de Biología Molecular y Medicina Regenerativa –CABIMER; Consejo Superior de Investigaciones Científicas; Universidad de Sevilla; Universidad Pablo de Olavide; Seville, Spain

**Keywords:** parental histone deposition, homologous recombination, translesion synthesis, DNA damage tolerance, replication fork, Mcm2, Dpb3

## Abstract

The advance and stability of replication forks rely on a tight co-regulation of the processes of DNA synthesis and nucleosome assembly. We have addressed the relevance of parental histone recycling in the mechanisms of DNA damage tolerance (DDT) – homologous recombination (HR) and translesion synthesis (TLS) – that assist replication forks under conditions that block their advance. We show that mutants affected in the deposition of parental histones are impaired in the recombinational repair of the single-strand DNA gaps generated during DDT, with the defects being more severe in mutants impaired in the lagging strand-specific deposition pathway. These recombinational defects are not due to a deficit of parental histones at the nascent strands but to an excess of parental nucleosomes at the invaded strand that destabilizes the sister chromatid junction formed after strand invasion. In conclusion, parental histone distribution at stressed forks regulates HR and provides a potential mechanism for the choice between HR and TLS that would depend on whether DNA synthesis is blocked at the lagging or the leading strand.

## Introduction

Chromosome duplication during S phase involves DNA replication and assembly into chromatin. These two processes can be challenged by multiple stress conditions that cause loss of DNA integrity and alterations in the pattern of nucleosome-associated epigenetic marks, which are linked to genetic disorders and cancer (Halazonetis et al. 2008; Jasencakova and Groth 2010). Chromatin assembly is physically coupled to and genetically co-regulated with DNA synthesis, leading to a rapid deposition of nucleosomes right behind the replication fork (Prado and Maya 2017). Among other functions, this timely supply of histones protects forks from collapsing (Clemente-Ruiz and Prado 2009; Clemente-Ruiz et al. 2011; Mejlvang et al. 2014). In addition, replication-coupled chromatin assembly facilitates the inheritance of epigenetic marks by randomly distributing between the sister chromatids the parental histones that are removed ahead of the fork. This source of histones is complemented with newly synthesized ones to completely package the duplicated DNA (Annunziato 2015). Therefore, cells have to coordinate a balanced deposition of old and new histones, each one labelled with different marks aimed to different functions.

A major challenge for cell is the presence of DNA lesions and adducts that hamper the advance of the replisome, as they threat the integrity of the replication fork and the timely completion of genome duplication. Cells are endowed with DNA damage tolerance (DDT) mechanisms that can be grouped in two according to the strategy employed to bypass the obstacle and fill in the stretches of single-strand DNA (ssDNA) generated as a consequence of the stalling of the fork. Translesion synthesis (TLS) mechanisms use low-fidelity polymerases to incorporate a nucleotide opposite the lesion, whereas homologous recombination (HR) mechanisms use the intact sister chromatid to circumvent the obstacle and fill in the ssDNA gap (Prado 2018; Sale et al. 2012). Although the contribution of the different TLS and HR mechanisms depends on the dose and type of blocking lesion, as well as the cell cycle stage and organism, a conserved feature from yeast to human is the regulatory role of the replication processivity factor PCNA. In response to an accumulation of ssDNA at the stalled fork, PCNA can be either monoubiquitylated at lysine 164 by the Rad6 (E2)-Rad18 (E3) complex to promote the recruitment of TLS polymerases or polyubiquitylated at the same residue by the Mms2-Ubc13 (E2)-Rad5 (yeast) / HLTF and SHPRH (mammal) (E3) complex to facilitate HR (Hoege et al. 2002; Davies et al. 2008; Daigaku et al. 2010; Niimi et al. 2008; Motegi et al. 2008). In mammal these HR events are mostly associated with fork reversal, where the nascent strands are displaced and reannealed to form a Holiday junction (HJ)-like structure (Zellweger et al. 2015; Krishnamoorthy et al. 2021). In contrast, most HR events in yeast are associated with the formation of post-replicative sister chromatid junctions (SCJs) (Liberi et al. 2005; Branzei et al. 2008), although a role for the helicase Mph1 in fork reversal has also been proposed (Choi et al. 2010; Zapatka et al. 2019).

PCNA sumoylation at lysines 127 and 164 by Ubc9-Siz1 (yeast) /RPC (mammal) introduces an additional level of control by recruiting the antirecombinogenic helicases Srs2 (yeast) and PARI (mammal) to prevent PCNA ubiquitylation (UbPCNA)-independent unscheduled recombinogenic rearrangements, which can be detected in mutants defective in both PCNA ubiquitylation and Srs2 (Schiestl et al. 1990; Hoege et al. 2002; Pfander et al. 2005; Papouli et al. 2005; Moldovan et al. 2012). As PCNA sumoylation is restricted to S phase, UbPCNA-independent HR is thought to operate in G2/M, in contrast to UbPCNA-dependent HR that operates mostly in S phase (Hoege et al. 2002; Karras and Jentsch 2010; Pfander et al. 2005; Papouli et al. 2005; Daigaku et al. 2010; Ortiz-Bazán et al. 2014).

Given the tight connection between DNA synthesis and nucleosome assembly at the fork, it is not surprising to find key regulatory functions for histone deposition factors in the DDT response. An example is the acetylation of histone H3 at lysine 156 (H3K156), a post-translational modification of newly synthesized histones that facilitates their deposition at forks (Li et al. 2008). This modification is maintained at chromatin under replicative stress (Masumoto et al. 2005), where it seems to promote ubiquitylation of some unknown substrate to uncouple the replicative helicase from the polymerases as a prerequisite for HR to bypass the blocking lesion (Wurtele et al. 2011; Luke et al. 2006; Buser et al. 2016; Zaidi et al. 2008; Luciano et al. 2015). In human cells, where H3K56 is a minor modification, a similar role has been reported for the MMS22L-TONSL complex, which binds to newly synthesized H3-H4K20me0 (unmethylated at lysine 20) at nascent chromatin to recruit and stabilize the recombinase RAD51 (Saredi et al. 2016).

Two specific and conserved pathways transfer parental H3-H4 tetramers to nascent chromatids: 1) the Dpb3 and Dpb4 subunits of the leading-strand polymerase (Pole) (POLE3 and POL4 in mammal) transfer parental histones to the leading strand, and 2) the axis formed by the Mcm2 subunit of the MCM helicase, Ctf4 and the polymerase a (Pol a) transfer parental histones to the lagging strand (Yu et al. 2018; Li et al. 2020; Gan et al. 2018; Foltman et al. 2013; Bellelli et al. 2018; Huang et al. 2015; Richet et al. 2015; Petryk et al. 2018). In this study we took advantage of specific mutants affected in either the lagging or the leading-specific parental histone deposition pathways to directly address their role in the DDT response. Our results show that proper parental histone recycling is required for efficient recombinational filling of ssDNA gaps during DDT. Specifically, an excess of parental nucleosomes at the invaded strand as a consequence of unbalance distribution of parental histones between the nascent chromatids destabilizes SCJs. Remarkably, these defects are more severe when the lagging-specific parental histone deposition pathway is impaired, which suggests a separation of labor for HR and TLS at the lagging and leading strands, respectively.

## Results

### Parental histone recycling contributes to MMS resistance through HR

To address a putative role of parental histone recycling in DDT we first analysed the sensitivity to methyl methanesulfonate (MMS) of mutants affected in the deposition of parental histones to the lagging *(mcm2-3A)* or the leading *(dpb3Δ)* strand (Gan et al. 2018; Foltman et al. 2013; Yu et al. 2018). Neither the single *mcm2-3A* and *dpb3Δ* mutants nor the double *mcm2-3A dpb3Δ* mutant displayed significant MMS sensitivity (Figure 1A). Accordingly, Rad53 was efficiently dephosphorylated after MMS treatment (Figure S1A). Only the *mcm2-3A* mutant showed a slight sensitivity at high MMS concentrations that was not further increased by a *pol1-2A* mutation (Figure 1B), consistent with Mcm2 and Pola operating in the same histone deposition pathway (Li et al. 2020; Gan et al. 2018). On the contrary, the *mcm2-3A* mutation conferred MMS sensitivity in combination with a partial reduction in the amount of Spt16 (Figure 1C), which is involved in the deposition of new and parental histones (Foltman et al. 2013; Yang et al. 2016). The contribution of parental histone recycling to MMS resistance became more evident when *mcm2-3A*, *pol1-2A* and *dpb3Δ* where combined with a leaky *rad5-535* allele, and this sensitivity was suppressed by expressing an additional copy of Spt16 in the *mcm2-3A rad5-535* mutant (Figure 1A). In accordance with these results, defective histone recycling delayed the checkpoint recovery after MMS treatment in cells expressing the *rad5-535* allele (Figure S1B). The MMS sensitivity and checkpoint recovery of the triple mutant *mcm2-3A dpb3Δ rad5-535* were similar to those displayed by the double mutants *mcm2-3A rad5-535* and *dpb3Δ rad5-535* (Figures 1A and S1B). This result was unexpected because a concomitant defect in the deposition of parental histones at the lagging and leading strands in the double mutant *mcm2-3A dpb3Δ* caused a synergistic alteration in nucleosome positioning (Schlissel and Rine 2019).

**Figure 1.**
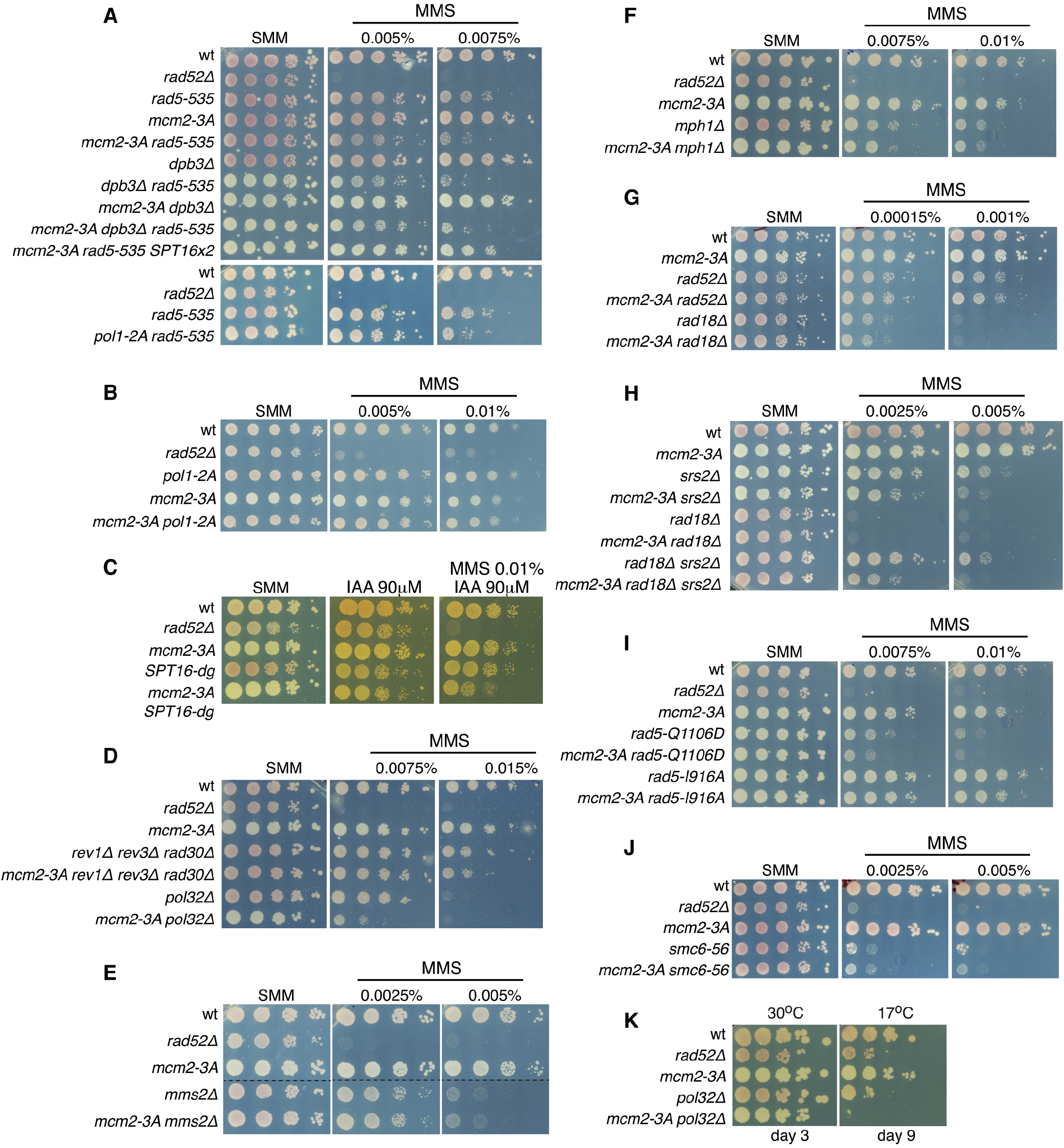
Parental histone recycling contributes to MMS resistance through HR. **(A-K)** Epistasis analyses between parental histone recycling and DDT mutants. MMS sensitivity of the indicated strains as determined by spotting ten-fold serial dilutions of the same number of midlog growing cells onto medium with or without the drug. The analyses were repeated at least twice with similar results.

To define what pathway of DDT was impaired in the histone recycling mutants, we analyzed epistatic relationships between *mcm2-3A* and mutants defective in TLS *(rev1Δ rev3Δ rad30Δ),* UbPCNA-dependent HR *(mms2Δ),* TLS and UbPCNA-dependent HR *(rad18Δ),* fork reversal *(mphlΔ’)* and HR *(rad52Δ)* (Figures 1D-G). The *mcm2-3A* allele only increased (slightly and at high doses) the MMS sensitivity of the *rev1Δ rev3Δ rad30Δ* mutant (Figure 1D), suggesting that parental histone deposition facilitates MMS resistance through a HR mechanism. Accordingly, the *mcm2-3A* allele increased the MMS sensitivity of a *rad18Δ srs2Δ* mutant (Figure 1H), in which UbPCNA-independent HR is the only operative DDT mechanism. Indeed, a similar additive effect on MMS sensitivity was observed in combination with the absence of Srs2 (Figure 1H).

The effect of *mcm2-3A* on MMS sensitivity was much stronger in cells expressing the *rad5-535* allele than lacking the TLS polymerases, suggesting that the synergistic MMS sensitivity with *rad5-535* is not related to the role of Rad5 in TLS (Gangavarapu et al. 2006; Gallo et al. 2019). The main function of Rad5 in DDT is associated with its ubiquitin ligase and helicase activities. These activities are specifically eliminated in *rad5-I916A* (ubiquitin ligase) and *rad5-Q1106D* (helicase) mutants (Ulrich 2003; Choi et al. 2015). The *mcm2-3A* mutation enhanced MMS sensitivity in both backgrounds (Figure 1I), suggesting that parental histone recycling facilitates both activities or an additional function.

Mutants affected in HR can partially rescue the MMS sensitivity of a *smc6-56* mutant impaired in SCJ resolution (Choi et al. 2015). Likewise, HR defective mutants can supress the cold sensitivity associated with the loss of Pol32 (Karras and Jentsch 2010), a subunit of both the TLS polymerase Pol ζ and the leading-associated polymerase Pol δ (Gerik et al. 1998; Johnson et al. 2012). The *mcm2-3A* mutation did not suppress any of these phenotypes (Figures 1J-K); actually, it further exacerbated both the cold sensitivity and the MMS sensitivity of the *pol32Δ* strain (Figures 1D and 1K). Since *mcm2-3A* hardly increased the MMS sensitivity of TLS mutants (Figure 1D), this synergistic effect might be associated with the loss of the Pol32-dependent function of Pol δ in MMS-induced HR, where it extents DNA synthesis upon strand invasion (Vanoli et al. 2010). However, although Pol32 is not essential for the replicative function of Pol δ (Gerik et al. 1998), its absence might destabilize lagging strand synthesis and exacerbate the histone deposition defects of the *mcm2-3A* mutant. In conclusion, our epistatic analyses suggest a function for parental histone recycling in MMS resistance that becomes more critical in combination with mutations affecting specific HR steps.

### Biased parental histone distribution impairs ssDNA gap filling by HR

As a first approach to understand the role of parental histone recycling in the DDT response, we analyzed the effect of *mcm2-3A* and *dpb3Δ* in the filling of ssDNA gaps generated during the replication of MMS-damaged DNA. The repair of these lesions occurs at centers that can be followed with GFP-labelled DNA repair proteins. For these experiments cells were synchronized in G1, released into S phase in the presence of 0.033% MMS for 1 hour and then into fresh medium after MMS inactivation to allow ssDNA filling. First, we followed the RPA complex (formed by Rfa1, 2 and 3 in *S. cerevisiae),* which covers ssDNA regardless of the mechanism of ssDNA filling. Moreover, the detection level of the Rfa1 signal is higher than that displayed by other repair proteins. Thus, it is an excellent and sensitive readout for the accumulation of unrepaired ssDNA fragments (Cano-Linares et al. 2021). Most wild type and histone deposition cells (~95%) accumulated RPA foci 1 hour after MMS inactivation (Figures 2A, left panel, and S2A). The percentage of wild type cells with foci dropped to 21% at the end of the time course (79% foci resolution), whereas it dropped to 61% and 37% in *mcm2-3A* and *dpb3Δ* cells (37% and 63% foci resolution, respectively) (Figure 2A, left and middle panels). Remarkably, the percentage of cells with Rad52 foci dropped to 43% in the double mutant *mcm2-3A dpb3Δ* (56% foci resolution). To get a more accurate calculation of the amount of unresolved ssDNA gaps, we determined the fluorescence signal at foci per cell, which integrates the percentage of cells that retain foci as well as the number and intensity of foci (Figure 2A, right panel). These analyses confirmed that the lagging-specific (*mcm2-3A*) but not the leading-specific (*dpb3Δ*) histone deposition mutant was severely affected in the filling of MMS-induced ssDNA gaps, and that this defect was partially suppressed by impairing histone deposition at the leading strand.

**Figure 2.**
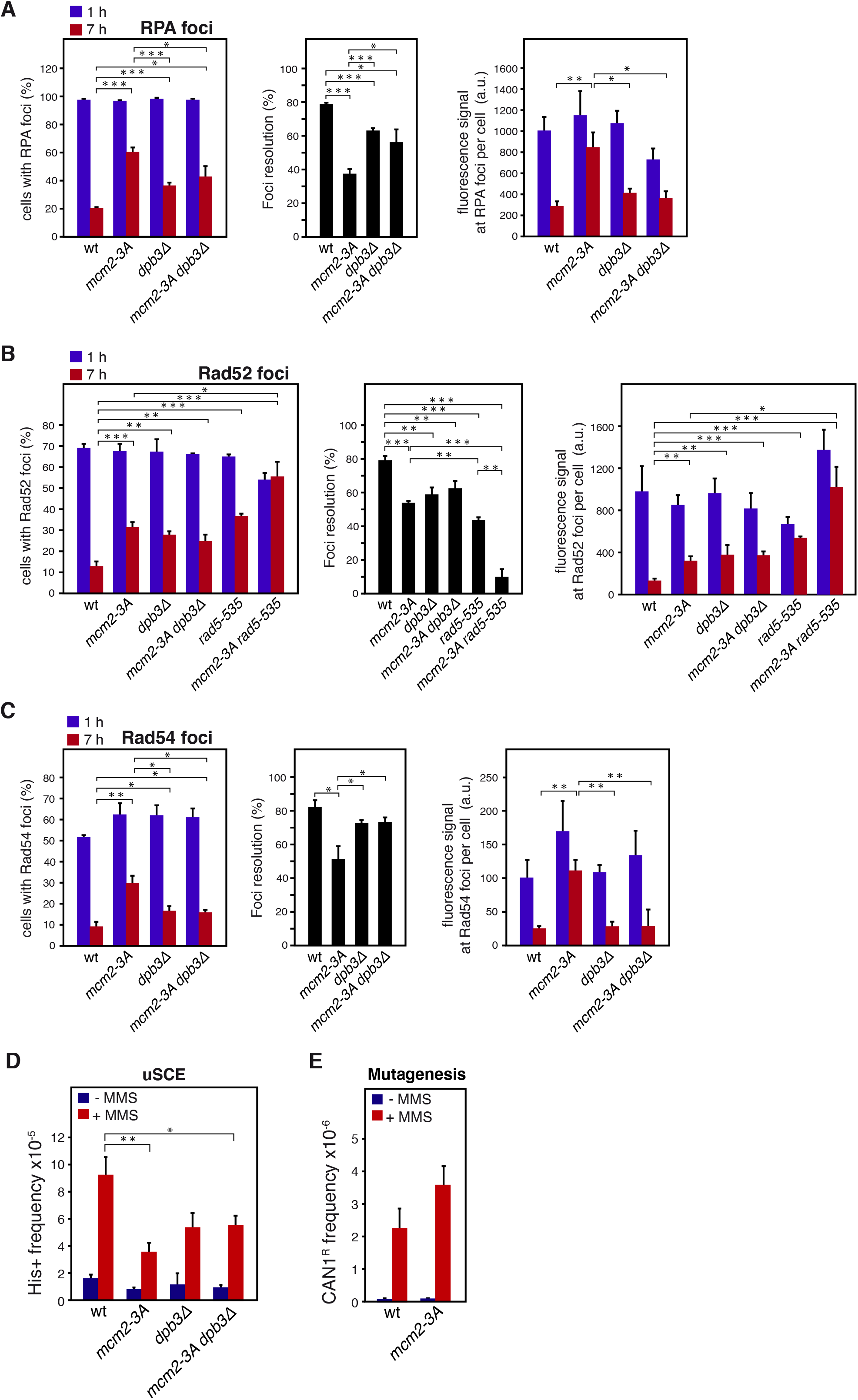
Biased parental histone distribution impairs ssDNA gap filling by HR. **(A-C)** DNA repair efficiency of parental histone recycling mutants and wild type cells, as determined by analysing the formation and resolution of MMS-induced Rfa1-YFP (A), Rad52-YFP (B) and Rad54-YFP (C) foci. Cells were synchronized in G1, released into S phase in the presence of 0.033% MMS for 1 hour and then into fresh medium after MMS inactivation. The percentage of cells with foci (left panels) and the fluorescent signal at foci per cell (right panels) at 1 (peak) and 7 hours after MMS treatment was determined with MetaMorph software. Foci resolution (middle panels) was calculated as (100 – percentage of cells with foci at 7 hours x 100)/ percentage of cells with foci at 1 hour. Complete kinetics and SuperPlots of the fluorescent signal at foci per cell are shown in Figure S2A-C. **(D-E)** Effect of parental histone recycling mutants in MMS-induced uSCE (D) and mutagenesis (E). The frequency of recombinants was determined from colonies grown in the absence and presence of 0.01% MMS. The means + SEM of 3 independent experiments (A-C) and 8-9 (D) or 3 (E) independent fluctuation tests are shown. Statistically significant differences according to an unpaired two-tailed Student’s *t-* test are shown, where one, two and three asterisks represent *P*-values <0.05, <0.01 and <0.001, respectively.

To define the ssDNA filling mechanism affected in these histone recycling mutants we followed the accumulation of the HR protein Rad52 at DNA repair centers. The efficiency of ssDNA gap filling, inferred from both the percentage of cells with foci and the Rad52 signal per cell at the end of the time course relative to the peak during the time course, was compromised in the histone recycling mutants *mcm2-3A* and *dpb3Δ*. However, the defect was slightly less severe in *dpb3Δ*, and importantly, epistatic over *mcm2-3A* (Figure 2B). Apart from its essential role in HR, Rad52 facilitates TLS (Cano-Linares et al. 2021). Therefore, we repeated the kinetics in cells expressing Rad54-YFP, which specifically labels HR-mediated gap filling events during DDT (Cano-Linares et al. 2021). In this case, the ssDNA gap filling defect was much more pronounced in *mcm2-3A* than in *dpb3Δ*, and again defective histone deposition at the leading strand partially suppressed the HR defect associated with defective histone deposition at the lagging strand (Figure 2C, compare *mcm2-3A dpb3Δ* with *mcm2-3A* and *dpb3Δ*). Finally, and in accordance with the MMS sensitivity and checkpoint recovery results, the double mutant *mcm2-3A rad5-535* displayed a synergistic defect in the resolution of MMS-induced Rad52 foci as compared to the single mutants (Figure 2B).

To determine if the defects in ssDNA gap filling were associated with a drop in the frequency of MMS-induced recombination events, we used an unequal sister chromatid exchange (uSCE) system where recombinants can be selected as His+ cells (Fasullo and Davis 1987). In line with the DNA repair foci results, impaired deposition of parental histones in the *mcm2-3A* and *dpb3Δ* mutants reduced the frequency of MMS-induced recombinants as compared to the wild type, this frequency was lower in *mcm2-3A* than in *dpb3Δ*, and the double mutant *mcm2-3A dpb3Δ* displayed the same frequency as the *dpb3Δ* mutant (Figure 2D). As expected, this defect in HR in the *mcm2-3A* mutant was accompanied by a slight increase in the alternative TLS pathway as inferred from the analysis of mutagenesis at the *CAN1* locus (Figure 2E). Altogether, these results indicate that parental histone recycling facilitates HR during DDT. This function is particularly sensitive to defects in the deposition of parental histones at the lagging strand, but these defects can be partially alleviated if both the leading and the lagging-specific histone deposition pathways are impaired. Therefore, efficient HR during DDT relies on a proper distribution of parental and newly synthesized histones on the sister chromatids.

### An excess of parental histones at the leading strand destabilizes HR-induced SCJs

To understand why a biased distribution of parental histones impairs HR we first analyzed the binding of the essential HR protein Rad52 to ssDNA gaps in the *mcm2-3A* mutant by chromatin-endogenous cleavage (ChEC). In this approach DNA binding is inferred from the amount of DNA digested by a chimera of Rad52 fused to MNaseI (Rad52-MN), whose nuclease domain is activated with Ca^2+^ ions (González-Prieto et al. 2013, 2021). MMS-induced DNA digestion by Rad52-MN was similar in *mcm2-3A* and wild type cells, suggesting that Rad52 binding to ssDNA gaps is not affected by impairing parental histone recycling (Figure 3A).

**Figure 3.**
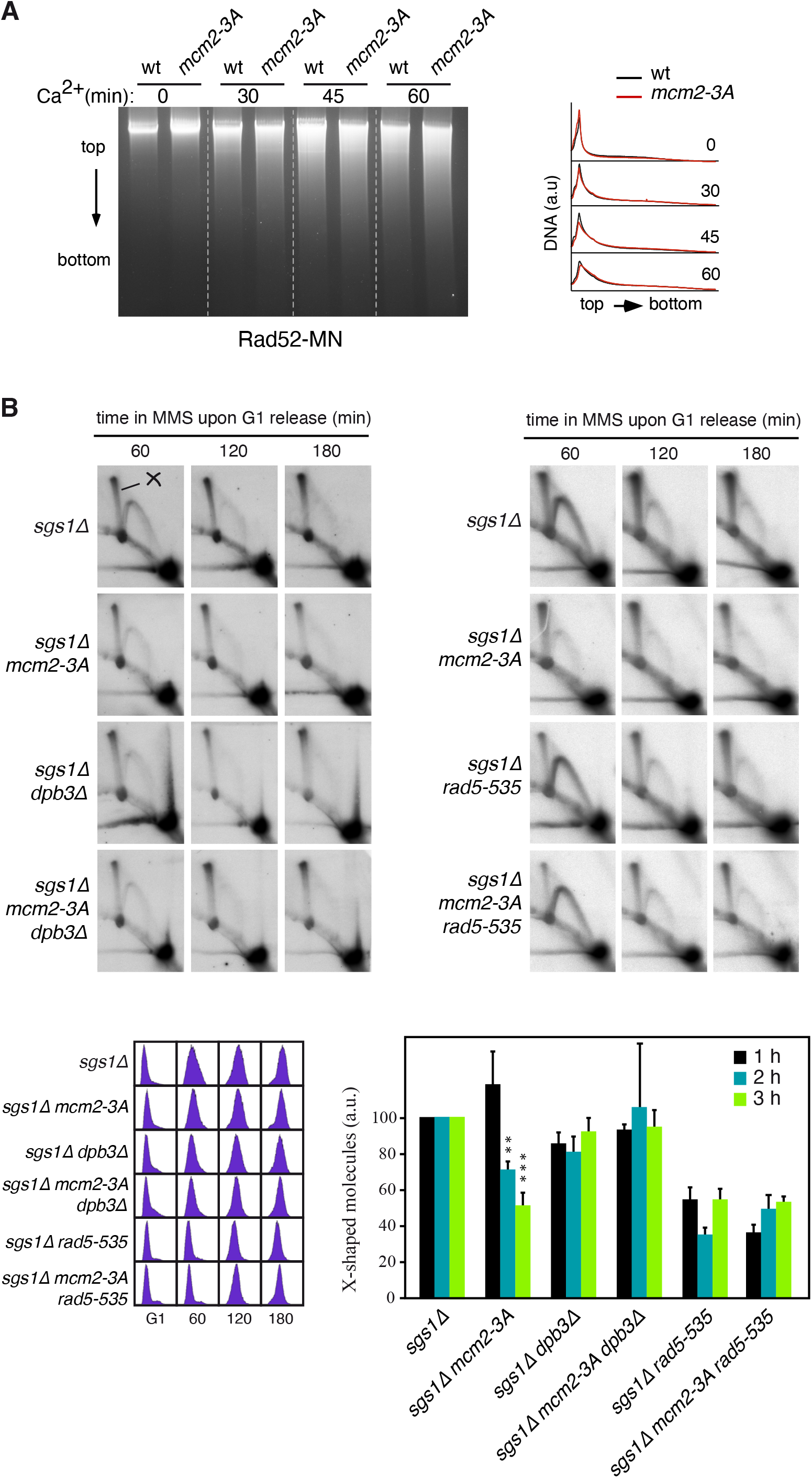
An excess of parental histones at the leading strand destabilizes HR-induced SCJs. **(A)** Rad52 binding to replicative ssDNA lesions in *mcm2-3A* and wild type cells, as determined by ChEC analysis of exponentially growing cells incubated with 0.05% MMS for 2 hours. Total DNA and DNA digestion profiles from cells permeabilized and treated with Ca^2+^ for different times are shown. The experiment was repeated twice with similar results. **(B)** SCJ formation in histone recycling mutants, as determined by 2D gel analysis of X-shaped molecules in cells synchronized in G1 and released in the presence of 0.033% MMS. The amount of X-shaped molecules (spike) relative to the total amount of molecules, with the wild type values set at 100, is shown. The mean + SEM of six (wild type and *mcm2-3A)* and three (rest) independent kinetics are shown. Statistically significant differences with the wild type (taken as 100) according to a two-tailed One sample *t*-test are shown, where two and three asterisks represent *P*-values <0.01 and <0.001, respectively.

The repair of MMS-induced ssDNA gaps by HR is associated with the formation of SCJs, which can be detected as X-shaped structures by 2D electrophoresis in *sgs1Δ* cells defective in the dissolution of the double HJ structure formed after invasion of the sister chromatid (Liberi et al. 2005). We asked if these recombination intermediates were affected in histone recycling mutants. Cells were synchronized in G1 and released into S phase for long times in the presence of 0.033% MMS, and the amount of SCJs was calculated relative to total molecules (Figure 3B). Defective parental histone deposition at the leading strand in the *dpb3Δ* mutant caused a slight but not significant reduction in the amount to X-shaped molecules relative to the wild type strain. On the contrary, the lagging-specific histone deposition mutant *mcm2-3A* displayed a slight increase at the beginning of the time course followed by a significant drop along the rest of the time course in X-shaped molecules. Therefore, SCJs form but are unstable when the deposition of parental histones at the lagging strand is impaired. Importantly, this defect was suppressed in the double mutant *mcm2-3A dpb3Δ* (Figure 3B), indicating that SCJ instability in the *mcm2-3A* mutant is not caused by a deficit of parental histones at the lagging strand but by an excess of parental histones at the leading strand. Finally, we repeated the analysis in *rad5-535* background. This mutation compromised the formation of SCJs; however, this defect was not exacerbated by the *mcm2-3A* allele (Figure 3B). Therefore, the additive effects of *rad5-535* and *mcm2-3A* in MMS sensitivity, checkpoint recovery and ssDNA gap filling are likely due to an additional function of parental histone deposition on DDT.

## Discussion

DDT is triggered by conditions that impair the advance of replication forks. Given the genetic and physical connection between DNA synthesis and histone deposition at forks, we decided to address the relevance of the parental histone deposition pathways in the mechanisms of DDT. We reveal the importance of parental histone recycling in the efficiency of ssDNA gap filling by HR during DDT, as determined by the analyses of mutants specifically affected in the deposition of parental histones at the lagging (*mcm2-3A*) and leading (*dpb3Δ)* strands. Our epistasis and molecular analyses show that these mutants are defective in the accumulation of SCJs, in line with a parallel work by Branzei and colleagues (Dolce et al. 2022). Our study extends this finding by revealing that the HR defects are not due to a loss of parental nucleosomes at the nascent strands but to a biased distribution of parental and newly synthesized histones. Specifically, the ssDNA gap filling and SCJ stability defects associated with impaired histone deposition at the lagging strand in the *mcm2-3A* mutant are partially suppressed by also affecting parental histone deposition at the leading strand in the double mutant *mcm2-3A dpb3Δ* (Figures 2 and 3). This implies that HR in the *mcm2-3A* mutant is not impaired by a deficit of parental histones at the lagging strand but by an excess of parental histones at the leading strand.

Excess parental histones at the leading strand of the *mcm2-3A* mutant might prevent HR at different molecular steps. We have discarded a role in the loading of the mediator protein Rad52 at MMS-induced ssDNA gaps. Actually, the analysis of HR intermediates showed that SCJs form at similar levels in *mcm2-3A* and wild type cells (Figure 3), indicating that Rad51-mediated strand-invasion and strand-exchange steps are not affected by altering the distribution of parental histones between the nascent strands. However, these SCJ intermediates are unstable and this instability is due to an excess of parental histones at the leading strand, as inferred by its suppression in the double mutant *mcm2-3A dpb3Δ.* Since this analysis was performed in cells defective in SCJ dissolution *(sgs1Δ* background), the excess of parental histones may destabilize the D-loop structure generated after strand invasion. Therefore, the excess of parental histones at the leading strand of the *mcm2-3A* mutant prevents the completion of those HR events that are initiated at the lagging strand and invade the leading strand (Figure 4; only nucleosomes at the invaded strand are shown for simplicity).

**Figure 4.**
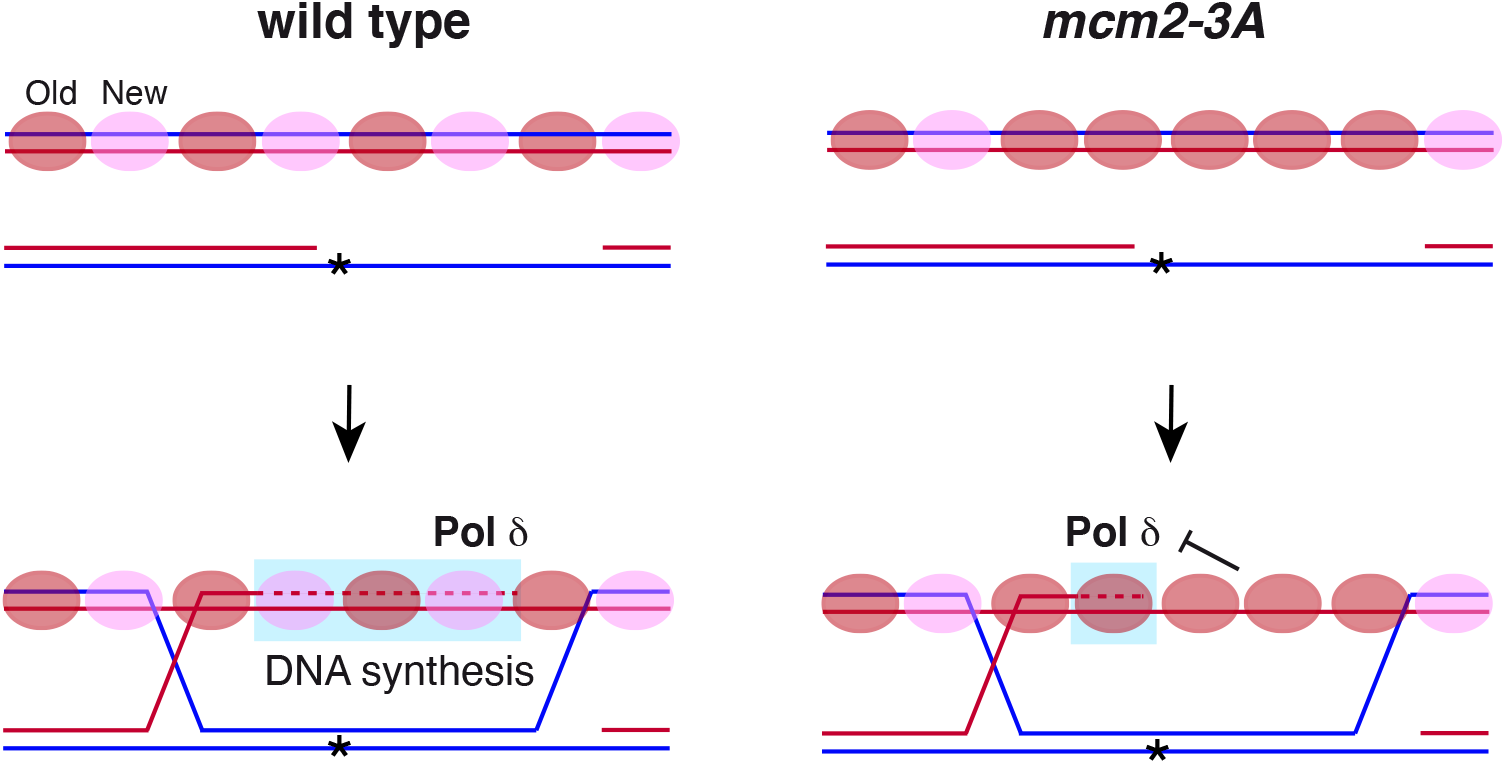
Parental histone distribution at stressed forks regulates HR during DDT. The recombinational filling of ssDNA gaps during DDT requires invasion of the intact sister chromatid and DNA synthesis at the 3’-invading end by Polδ. In a *mcm2-3A* mutant an excess of parental histones at the leading strand allows strand invasion but destabilizes SCJs before the formation of the double-HJ structure, suggesting that it is preferentially affecting completion of HR events that initiate at the lagging strand and invade the leading strand (only nucleosomes at the invaded strand are shown for simplicity). Excess parental histones might cause SCJ destabilization by either promoting some antirecombinogenic activity (e.g., Srs2) or counteracting the DNA synthesis activity of Polδ due to the higher stability of parental nucleosomes. See text for the biological implications of this model in the choice of the DDT mechanism depending on the position of the blocking lesion.

How an accumulation of parental histones at the invaded strand destabilizes SCJs is currently unknown. One possibility is that the accumulation of parental histones at the invaded strand augments the antirecombinogenic activity of Srs2. It has been shown in *Schizosaccharomyces pombe* that the assembly of newly synthesized histones at recombination intermediates by CAF1 and Asf1 facilitates HR-mediated fork restart by counteracting the D-loop disassembly activity of the DNA helicase Rqh1 (Pietrobon et al. 2014; Hardy et al. 2019). Likewise, the deficit of new histones that accompanies an excess of parental histones at the same strand (Gan et al. 2018; Yu et al. 2018) might partially remove histone post-translational modifications required to counteract the anti-recombinogenic activity of Srs2. Srs2 has been proposed to negatively control HR during DDT independently of its DNA helicase activity; instead, Srs2 seems to inhibit HR by promoting the disassembly of the PCNA/Polδ complex and the subsequent DNA synthesis-mediated D-loop extension (Burkovics et al. 2013). Inhibition of this Srs2 activity would better explain the formation but further instability of the SCJs. However, *mcm2-3A* increases the MMS sensitivity of *srs2Δ* cells (Figure 1H), suggesting that it impairs HR independently of Srs2. An alternative possibility is related to the higher stability of parental as compared with newly synthesized nucleosomes (Yu et al. 2018). In this frame, a high density of parental nucleosomes might hamper the advance of Polδ to copy the intact information in the invaded strand (Figure 4). Since DNA synthesis at the invaded strand beyond the sequence corresponding to the blocking lesion is essential for the recombinational repair of the ssDNA gap, a defect at this step might signal the disassembly of the SCJ. This hypothesis would explain the high MMS and cold sensitivity displayed by *mcm2-3A* in the absence of Pol32 (Figures 1D and 1K), which is required for Polδ-mediated DNA synthesis and SCJ stability during DDT (Vanoli et al. 2010). Nevertheless, we cannot rule out the possibility that these synergistic effects were a consequence of a more dramatic histone deposition problem due to impaired DNA processivity at the lagging strand, as shown in human cells where PARP inhibition interferes with both DNA synthesis and parental histone deposition at the lagging strand (Zwinderman et al. 2021).

It must be stressed that SCJ instability can explain some but not all DDT defects displayed by the histone recycling mutants because *rad5-535* was epistatic over *mcm2-3A* for the loss of SCJs. Therefore, the synergistic defects of these two mutations in MMS sensitivity, checkpoint recovery and ssDNA gap filling must be due to an additional and not-yet identified function of parental histone recycling on DDT. The lack of this function may also explain the increased MMS sensitivity observed after combining *rad5-535* and *dpb3Δ* mutations.

In human cells parental histones display a slight preference for the leading strand under unperturbed conditions, and this pattern is strongly inverted under replication stress conditions (e.g., dNTPs depletion, checkpoint inactivation). Since these conditions have a major impact in the processivity of the leading strand, this suggests that parental histone deposition is particularly sensitive to problems that uncouple the helicase and polymerase activities (Zwinderman et al. 2021). DNA adducts preferentially block the leading strand because a block in the lagging strand only inhibits DNA synthesis of the corresponding Okazaki fragment (Lopes et al. 2006; Hashimoto et al. 2011; Pagès and Fuchs 2003; Higuchi et al. 2003; McInerney and O’Donnell 2004). Thus, the major effect of MMS treatment on histone distribution is expected to be an accumulation of parental histones at the lagging strand of replisomes blocked at the leading, with this effect being attenuated in *mcm2-3A* and exacerbated in *dpb3Δ.* Since an accumulation of parental histones at the invaded strand impairs SCJ stability, it is particularly intriguing that the HR defects were more severe in the lagging-specific (*mcm2-3A*) than in the leading-specific (*dpb3Δ*) histone deposition mutant because they both display similar alterations in the distribution of parental histones (Gan et al. 2018; Yu et al. 2018). A possible explanation to this apparent paradox is that HR deals preferentially with blocks at the lagging strand, whereas TLS does it with blocks at the leading strand. This division of labor would be favoured by the higher processivity of the lagging relative to the leading strand under replication stress conditions (Lopes et al. 2006; Hashimoto et al. 2011; Gan et al. 2017; Yu et al. 2014), which will promote an asymmetric accumulation of parental histones that may bias the recombinational repair to the lagging strand. Furthermore, PCNA is unloaded from lagging strands and reloaded at leading strands under replication stress conditions (including MMS treatment), which might help to recruit TLS polymerases to the leading strand (Yu et al. 2014). This division of labor would have two important functional implications. First, it would facilitate DNA replication because recombination initiation at the leading, but not at the lagging, requires the disassembly of at least part of the replisome to either promote fork reversal or prime new DNA synthesis downstream the blocking lesion for a post-replicative recombinational filling of the ssDNA gap. Second, cells might control the accumulation of mutations (mostly associated with TLS) at specific regions during asymmetric cell division or development by regulating the histone deposition pathways. Future experiments will be required to validate this hypothesis and test if it is behind the differences in Rad54 and Rad52 foci resolution (impaired only in HR or in HR and TLS mutants, respectively (Cano-Linares et al. 2021) observed between *mcm2-3A* and *dpb3Δ* mutants. In conclusion, parental histone distribution at stressed forks regulates HR and provides a potential mechanism for the choice of the DDT mechanism.

## Supporting information

Gonzalez-Garrido.ms

## ACKNOWLEDGMENTS

We thank Karim Labib, Helle D. Ulrich and Xiaolan Zhao for various strains. This publication is part of the PGC2018-099182-B-100 grant, funded by MCIN/AEI/10.13039/501100011033 and FEDER “Una manera de hacer Europa”, and the P20_00750 grant, funded by the Andalusian government. CG-G is supported by a grant (FPU18/05207) from the MEFP (Spanish Government).

## AUTHOR CONTRIBUTION

Investigation, CG-G; conceptualization, FP; Writing original draft, FP; Writing, review and editing, FP; funding acquisition, FP.

## MATERIALS AND METHODS

### Yeast strains, plasmids and growth conditions

All *Saccharomyces cerevisiae* strains used are haploid derived from W303 (Table S1). Tagged and deletion strains were constructed by a PCR-based strategy (Longtine et al. 1998). The original uSCE strain (Fasullo and Davis 1987) was backcrossed seven times into the W303 background. The Spt16^dg^ strain was generated by tagging Spt16 at the carboxy-terminal end with AID following a “one-step PCR” approach in a strain expressing *Os*TIR under control of a constitutive *ADH1* promoter (Nishimura et al. 2009). Spt16 was degraded after 90 minutes in the presence of 500 μM indole-3-acetic acid (IAA) (data not shown). pHyg-AID-9myc is a plasmid for AID tagging (Morawska and Ulrich 2013) pWJ1213 is a centromeric plasmid expressing *RAD52-YFP* (Alvaro et al. 2006). Yeast cells were grown at 30°C in supplemented minimal medium (SMM). For G1 synchronization, cells were grown to mid-log phase and a-factor was added twice at 60 min intervals at either 2 *(BAR1* strains) or 0.25 μg/ml (*bar1Δ* strains). Then, cells were washed three times and released into fresh medium with 50 μg/ml pronase in the absence or presence of MMS at the indicated concentrations. To eliminate the MMS before releasing cells into fresh medium, samples were treated with 2.5% sodium thiosulfate to inactivate it and then washed three times. MMS sensitivity was determined by spotting ten-fold serial dilutions of the same number of mid-log growing cells onto medium with or without the drug.

### Flow cytometry

DNA content analysis was performed by flow cytometry as reported previously (Prado and Aguilera 2005). Cells were fixed with 70% ethanol, washed with phosphate-buffered saline (PBS), incubated with 1 mg of RNaseA/ml PBS, and stained with 5 μg/ml propidium iodide. Samples were sonicated to separate single cells and analysed in a FACSCalibur flow cytometer.

### Genetic recombination and mutagenesis assays

HR was determined by measuring the frequency of His+ recombinants generated by uSCE in a chromosomal-integrated system (Fasullo and Davis 1987), whereas mutagenesis was determined by measuring the frequency of forward mutagenesis at the *CAN1* locus (selected as canavanine-resistant cells). Recombination and mutant frequencies were determined by fluctuation tests as previously reported (Cano-Linares et al. 2021). Briefly, cells from six independent colonies of similar size and isolated on medium with +/- MMS were plated with the appropriate dilutions onto SMM without histidine, SMM without arginine but containing 60 μg/ml canavanine and SMM to calculate recombinants, mutants and total viable cells (as colony-forming units), respectively. The frequencies of HR and mutation were calculated using the median of recombinants/mutants and the mean of total cells. To have a more accurate value, the mean and SEM of at least 3 independent fluctuation tests are given.

### DNA repair foci analysis

The percentage of cells with RPA, Rad52 and Rad54 foci was determined as described previously (Cano-Linares et al. 2021). Cells expressing Rfa1-YFP, Rad54-YFP or transformed with plasmid pWJ1213 (expressing Rad52-YFP) were grown in liquid culture under the indicated conditions, fixed with 2.5% formaldehyde in 0.1M potassium phosphate pH 6.4 for 10 minutes, washed twice with 0.1M potassium phosphate pH 6.6 and resuspended in 0.1M potassium phosphate pH 7.4. Finally, cells were fixed with 80% ethanol for 10 minutes, resuspended in H2O and visualized with a Leica CTR6000 fluorescence microscope. The percentage of cells with foci was counted directly on the processed samples under the microscope or on acquired images. In this case, six contrast and fluorescence images along the z-axis (0.49 μm length each) were acquired with a CCD camera (Leica DFC350 FX) to find well-defined foci. Images were processed and analyzed with the MetaMorph software (Molecular devices). A total number of approximately 100 cells were analyzed for each time point and experiment.

### 2D-gel electrophoresis

Replication intermediates were analyzed by 2D-gel electrophoresis from cells arrested with sodium azide (0.1% final concentration) and cooled down on ice as reported (Clemente-Ruiz and Prado 2009). Briefly, total DNA was isolated with the G2/CTAB protocol, digested with specific restriction enzymes (*Eco*RV and *Hind*III), resolved by neutral/neutral two-dimensional gel electrophoresis, blotted to Hybond^TM^-XL membranes, and analyzed by hybridization with ^32^Plabelled probe A for the *ARS305* proximal fragment (Clemente-Ruiz and Prado 2009). All signals were acquired in a Fuji FLA5100 and quantified with the ImageGauge software (Fujifilm).

### *In vivo* ChEC analyses

Chromatin endogenous cleavage (ChEC) of Rad52-MN cells was performed as reported (González-Prieto et al. 2013, 2021) from asynchronous cultures grown in the presence of 0.05% MMS for 2 hours and arrested with sodium azide (0.1% final concentration). For cleavage induction, digitonin-permeabilized cells were incubated with 2 mM CaCl_2_ at 30°C under gentle agitation. Total DNA was isolated and resolved into 0.8% TAE l’ agarose gels. Gels were scanned in a Fuji FLA5100, and the signal profile quantified using the ImageGauge software (Fujifilm). The area of the DNA digestion profiles was equalized to eliminate DNA loading differences.

### Western blot

Yeast protein extracts were prepared using the TCA protocol (FOIANI et al. 1994), resolved by 8% SDS-PAGE, and probed with a rabbit polyclonal antibody against Rad53 (Abeam, abl04232). All western signals were acquired and quantified in a ChemiDoc MP image system and quantified with the Image Lab^TM^ software (Biorad).

### Statistical analyses

Statistical analyses were performed using the Prism software (GraphPad). Mean, SEM, sample size and statistical tests are indicated in the Figure legends. Sample size was not predetermined using statistical methods. Given the reduced sample size, the analyses were performed assuming that they follow normal distributions. Scatter SuperPlots were done as recently reported (Lord et al. 2020).

---

**Figure S1. Defective histone recycling delays the checkpoint recovery after MMS treatment in cells expressing the *rad5-535* allele.**

**(A)** Defective parental histone deposition in *mcm2-3A* and *dpb3Δ* mutants does not affect checkpoint recovery.

**(B)** Defective histone recycling in *mcm2-3A* and *dpb3Δ* mutants delays the checkpoint recovery after MMS treatment in cells expressing the *rad5-535* allele.

Cells were synchronized in G1, released into S phase in the presence of 0.033% MMS for 1 hour and then into fresh medium after MMS inactivation. Checkpoint activation and recovery was followed by western blot against Rad53 (MMS-induced Rad53 phosphorylation causes a mobility shift). The experiments were repeated twice with similar results.

**Figure S2. Figure supplementary to Figure 2.**

**(A-C)** Complete kinetics and SuperPlots of the fluorescent signal at foci per cell from experiments reported in Figures 2A-C. The percentage of cells with foci in the complete kinetics was counted directly on the processed samples under the microscope. Cell cycle profiles were determined by cell sorting.

