## Supplementary material for "Parental histone distribution at nascent strands controls homologous recombination during DNA damage tolerance": Gonzalez-Garrido.ms

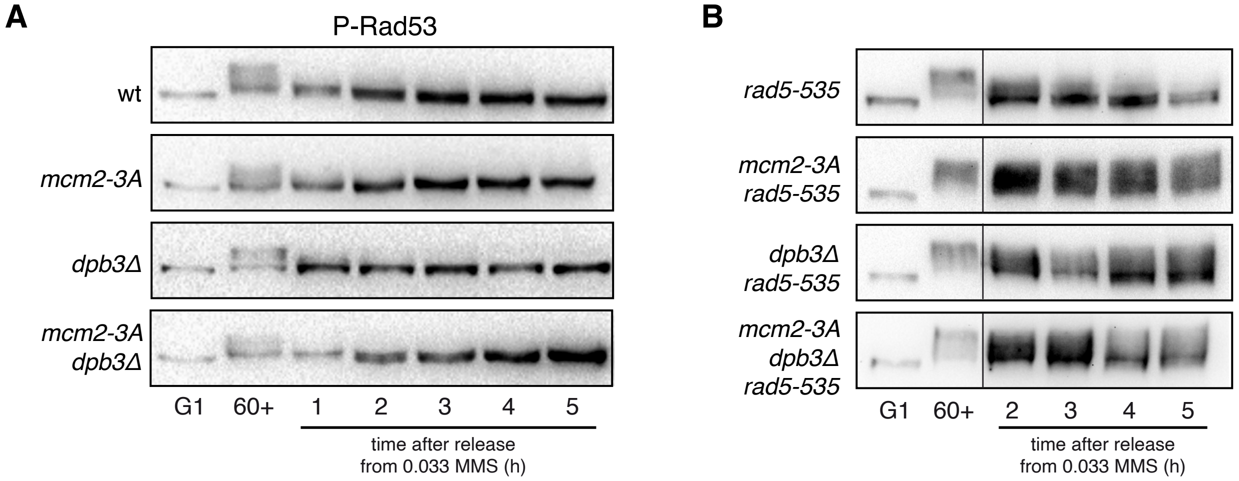
A
P-Rad53
wt
mcm2-3A
dpb3Δ
mcm2-3A
dpb3Δ
G1
60+
1
2
3
4
5
time after release
from 0.033 MMS (h)
B
rad5-535
mcm2-3A
rad5-535
dpb3Δ
rad5-535
mcm2-3A
dpb3Δ
rad5-535
G1
60+
2
3
4
5
time after release
from 0.033 MMS (h)

González-Garrido.Figure S2

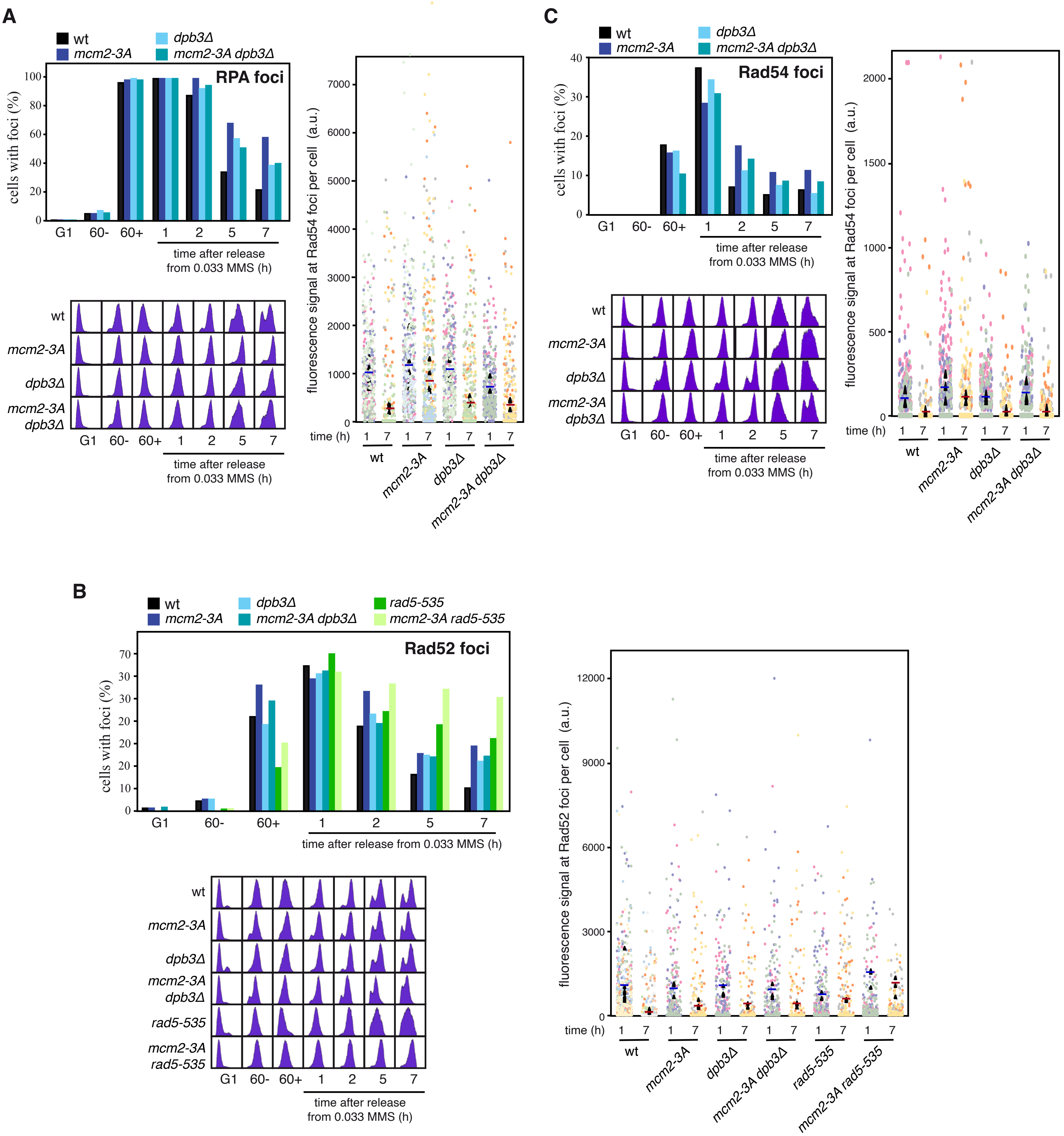

**Table S1.** ***Saccharomyces cerevisiae*** **strains used in this study**

| **Strain** | ***Genotype*** | **Ref.** | **Fig.** |
|---|---|---|---|
| W303-1aR5 | *MATa leu2-3,112 trp1-1 ura3-1 ade2-1 his3-11,15 can1-100 BAR1 RAD5* | (González-Prieto et al. 2013) | 1A-K, S1A, S2B |
| Wr52-5A | *MATa leu2-3,112 trp1-1 ura3-1 ade2-1 his3-11,15 can1-100 BAR1 rad52Δ::HYGMX4 RAD5* | (Barrientos-Moreno et al. 2018) | 1A-F,1I-K |
| Wm2-3A | *MATa leu2-3,112 trp1-1 ura3-1 ade2-1 his3-11,15 can1-100 BAR1 pep4Δ::ADE2 mcm2-3A (hphNT) RAD5* | This work | 1A-K,2B, S1A, S2B |
| Wd3 | *MATa leu2-3,112 trp1-1 ura3-1 ade2-1 his3-11,15 can1-100 BAR1 dpb3Δ::KANMX4 RAD5* | This work | 1A,2B, S1A, S2B |
| Wm2d3-3D | *MATa leu2-3,112 trp1-1 ura3-1 ade2-1 his3-11,15 can1-100 BAR1 pep4Δ::ADE2 mcm2-3A (hphNT) dpb3Δ::KANMX4 RAD5* | This work | 1A,2B, S1A, S2B |
| Wm2Ar5-4D | *MATa leu2-3,112 trp1-1 ura3-1 ade2-1 his3-11,15 can1-100 BAR1 mcm2-3A (hphNT) rad5-535* | This work | 1A,2B, S1B, S2B |
| Wd3r5-1B | *MATa leu2-3,112 trp1-1 ura3-1 ade2-1 his3-11,15 can1-100 BAR1 dpb3Δ::KANMX4 rad5-535* | This work | 1A, S1B |
| Wm2d3r5-7C | *MATa leu2-3,112 trp1-1 ura3-1 ade2-1 his3-11,15 can1-100 BAR1 pep4Δ::ADE2 mcm2-3A (hphNT) dpb3Δ::KANMX4 rad5-535* | This work | 1A, S1B |
| Wp1r5-3A | *MATa leu2-3,112 trp1-1 ura3-1 ade2-1 his3-11,15 can1-100 BAR1 pol1-2A2(hphNT) rad5-535* | This work | 1A |
| Wr5-2A | *MATa leu2-3,112 trp1-1 ura3-1 ade2-1 his3-11,15 can1-100 BAR1 rad5-535* | This work | 1A,2B, S1B, S2B |
| YMP584-2 | *MATa leu2-3,112 trp1-1 ura3-1 ade2-1 his3-11,15 can1-100 pep4Δ::ADE2 mcm2-3A (hphNT) leu2-3,112::pRS305-SPT16-9MYC (LEU2) rad5-535* | (Foltman et al. 2013) | 1A |
| Wp1-11C | *MATa leu2-3,112 trp1-1 ura3-1 ade2-1 his3-11,15 can1-100 BAR1 pol1-2A2 (hphNT) RAD5* | This work | 1B |
| Wm2-3Ap1 | *MATa leu2-3,112 trp1-1 ura3-1 ade2-1 his3-11,15 can1-100 BAR1 mcm2-3A (hphNT) pol1-2A2 (hphNT) RAD5* | This work | 1B |
| WSpt16$^{Dg}$-3C | *MATa leu2-3,112 trp1-1 ura3-1 ade2-1 his3-11,15 can1-100 ura3-1::ADH1-OsTIR1-9Myc(URA3) Spt16:: Spt16-aid (HygMX) RAD5* | This work | 1C |
| WSpt16$^{Dg}$ m2-13D | *MATα leu2-3,112 trp1-1 ura3-1 ade2-1 his3-11,15 can1-100 pep4Δ::ADE2 ura3-1::ADH1-OsTIR1-9Myc(URA3) Spt16:: Spt16-aid (HygMX) mcm2-3A(hphNT) RAD5* | This work | 1C |
| Wr1r3r30 | *MATa leu2-3,112 trp1-1 ura3-1 ade2-1 his3-11,15 can1-100 bar1Δ::LEU2 rev1Δ::NATMX4 rev3Δ::KANMX4 rad30::HYGMX4 RAD5* | (Cano-Linares et al. 2021) | 1D |

| | | | |
|---|---|---|---|
| Wr1r3r30m2-11A | *MATa leu2-3,112 trp1-1 ura3-1 ade2-1 his3-11,15 can1-100 bar1Δ::LEU2 rev1Δ::NATMX4 rev3Δ::KANMX4 rad30Δ::HYGMX4 mcm2-3A (hphNT) RAD5* | This work | 1D |
| Wpol32 | *MAT a leu2-3,112 trp1-1 ura3-1 ade2-1 his3-11,15 can1-100 BAR1 pol32Δ::HYGMX4 RAD5* | (Cano-Linares et al. 2021) | 1D, K |
| Wpl32m2-1D | *MATa leu2-3,112 trp1-1 ura3-1 ade2-1 his3-11,15 can1-100 BAR1 pep4Δ::ADE2 pol32 Δ::HYGMX4 mcm2-3A (hphNT) RAD5* | This work | 1D, K |
| Wmms2-2B | *MATa leu2-3,112 trp1-1 ura3-1 ade2-1 his3-11,15 can1-100 BAR1 mms2Δ::HIS3 RAD5* | This work | 1E |
| Wmms2m2-1C | *MATa leu2-3,112 trp1-1 ura3-1 ade2-1 his3-11,15 can1-100 BAR1 mms2Δ::HIS3 mcm2-3A (hphNT) RAD5* | This work | 1E |
| Wmph1 | *MATa leu2-3,112 trp1-1 ura3-1 ade2-1 his3-11,15 can1-100 BAR1 mph1Δ::KANMX4 RAD5* | This work | 1F |
| Wmph1m2 | *MATa leu2-3,112 trp1-1 ura3-1 ade2-1 his3-11,15 can1-100 BAR1 pep4Δ::ADE2 mph1Δ::KANMX4 mcm2-3A (hphNT) RAD5* | This work | 1F |
| Wr52Δm2-2B | *MATa leu2-3,112 trp1-1 ura3-1 ade2-1 his3-11,15 can1-100 BAR1 pep4Δ::ADE2 rad52Δ::HYGMX4 mcm2-3A (hphNT) RAD5* | This work | 1G |
| Wr18-2B | *MATa leu2-3,112 trp1-1 ura3-1 ade2-1 his3-11,15 can1-100 BAR1 rad18Δ::KANMX4 RAD5* | This work | 1G |
| Wr18m2-1C | *MATa leu2-3,112 trp1-1 ura3-1 ade2-1 his3-11,15 can1-100 BAR1 rad18Δ::KANMX4 mcm2-3A::HYGMX4 RAD5* | This work | 1G |
| Wsrs2-8B | *MATa leu2-3,112 trp1-1 ura3-1 ade2-1 his3-11,15 can1-100 BAR1 pep4Δ::ADE2 srs2Δ::KANMX4 RAD5* | This work | 1H |
| Wsrs2m2-15D | *MATa leu2-3,112 trp1-1 ura3-1 ade2-1 his3-11,15 can1-100 BAR1 pep4Δ::ADE2 srs2Δ::KANMX4 mcm2-3A (hphNT) RAD5* | This work | 1H |
| Wsrs2r18-3A | *MATa leu2-3,112 trp1-1 ura3-1 ade2-1 his3-11,15 can1-100 barΔ::LEU2 srs2Δ::KANMX4 rad18Δ::KANMX4 RAD5* | This work | 1H |
| Wsrs2r18m2-1D | *MATa leu2-3,112 trp1-1 ura3-1 ade2-1 his3-11,15 can1-100 BAR1 pep4Δ::ADE2 srs2Δ::KANMX4 rad18Δ::KANMX4 mcm2-3A (hphNT) RAD5* | This work | 1H |
| Wr5-I916A-1D | *MATa leu2-3,112 trp1-1 ura3-1 ade2-1 his3-11,15 can1-100 BAR1 rad5-I916A::KANMX4* | This work | 1I |
| Wr5-I916Am2-2D | *MATa leu2-3,112 trp1-1 ura3-1 ade2-1 his3-11,15 can1-100 BAR1 pep4Δ::ADE2 rad5-I916A::KANMX4 mcm2-3A (hphNT)* | This work | 1I |

| | | | |
|---|---|---|---|
| Wr5-Q1106D-2B | *MATa leu2-3,112 trp1-1 ura3-1 ade2-1 his3-11,15 can1-100 BAR1 pep4Δ::ADE2 rad5-Q1106D::KANMX4* | This work | 1I |
| Wr5-Q1106Dm2-1B | *MATa leu2-3,112 trp1-1 ura3-1 ade2-1 his3-11,15 can1-100 BAR1 pep4Δ::ADE2 rad5-Q1106-D::KANMX4 mcm2-3A (hphNT)* | This work | 1I |
| Wsmc6-56-4D | *MATa leu2-3,112 trp1-1 ura3-1 ade2-1 his3-11,15 can1-100 BAR1 smc6-56::KANMX4 RAD5* | This work | 1J |
| Wsmc6-56m2-3C | *MATa leu2-3,112 trp1-1 ura3-1 ade2-1 his3-11,15 can1-100 BAR1 smc6-56::KANMX4 mcm2-3A (hphNT) RAD5* | This work | 1J |
| WRFA1YFP | *MATa leu2-3,112 trp1-1 ura3-1 ade2-1 his3-11,15 can1-100 bar1Δ::LEU2 RFA1-8ala-YFP RAD5* | (Cano-Linares et al. 2021) | 2A, S2A |
| WRFA1YFPm2-7D | *MATa leu2-3,112 trp1-1 ura3-1 ade2-1 his3-11,15 can1-100 mcm2-3A (hphNT) RFA1-8ala-YFP RAD5* | This work | 2A, S2A |
| WRFA1YFPd3-5A | *MATa leu2-3,112 trp1-1 ura3-1 ade2-1 his3-11,15 can1-100 BAR1 dpb3Δ::KANMX4 RFA1-8ala-YFP RAD5* | This work | 2A, S2A |
| WRFA1YFPm2d3-1A | *MATa leu2-3,112 trp1-1 ura3-1 ade2-1 his3-11,15 can1-100 BAR1 mcm2-3A (hphNT) dpb3Δ::KANMX4 hml::LEU2 RAD5 RFA1-8ala-YFP RAD5* | This work | 2A, S2A |
| WRAD54YFP-2B | *MAT a leu2-3,112 trp1-1 ura3-1 ade2-1 his3-11,15 can1-100 BAR1 RAD54YFP::URA3 RAD5* | This work | 2C, S2C |
| WRAD54YFPm2-4D | *MAT a leu2-3,112 trp1-1 ura3-1 ade2-1 his3-11,15 can1-100 RAD54YFP::URA3 mcm2-3A (hphNT) RAD5* | This work | 2C, S2C |
| WRAD54YFPd3-11B | *MAT a leu2-3,112 trp1-1 ura3-1 ade2-1 his3-11,15 can1-100 RAD54YFP::URA3 dpb3Δ::KANMX4 RAD5* | This work | 2C, S2C |
| WRAD54YFPm2d3-4B | *MAT a leu2-3,112 trp1-1 ura3-1 ade2-1 his3-11,15 can1-100 RAD54YFP::URA3 dpb3Δ::KANMX4 mcm2-3A (hphNT) RAD5* | This work | 2C, S2C |
| WSCE-1A | *MATa leu2-3,112 ura3-1 ade2-1 his3-11,15 can1-100 trp1::SCE::URA3 RAD5* | This work | 2D |
| WSCEm2-7C | *MATa leu2-3,112 ura3-1 ade2-1 his3-11,15 can1-100 trp1::SCE::URA3 mcm2-3A (hphNT) RAD5* | This work | 2D |
| WSCEd3-4D | *MATa leu2-3,112 ura3-1 ade2-1 his3-11,15 can1-100 trp1::SCE::URA3 dpb3Δ::KANMX4 RAD5* | This work | 2D |
| WSCEm2d3-2D | *MATa leu2-3,112 ura3-1 ade2-1 his3-11,15 can1-100 trp1::SCE::URA3 can1-100 mcm2-3A (hphNT) RAD5* | This work | 2D |
| W303CAN | *MATa leu2-3,112 trp1-1 ura3-1 ade2-1 his3-11,15 BAR1 RAD5* | (Cano-Linares et al. 2021) | 2E |

| | | | |
|---|---|---|---|
| WCANm2-8B | *MATa leu2-3,112 trp1-1 ura3-1 ade2-1 his3-11,15 BAR1 mcm2-3A (hphNT) RAD5* | This work | 2E |
| WRad52MN-14B | *MATa leu2-3,112 trp1-1 ura3-1 ade2-1 his3-11,15 can1-100 barΔ::HisG Rad52MN::HISMX6 RAD5* | This work | 3A |
| WRad52MNm2-1A | *MATa leu2-3,112 trp1-1 ura3-1 ade2-1 his3-11,15 can1-100 barΔ::HisG Rad52MN::HISMX6 mcm2-3A (hphNT) RAD5* | This work | 3A |
| W303sgs1 | *MATa leu2-3,112 trp1-1 ura3-1 ade2-1 his3-11,15 can1-100 BAR1 sgs1Δ::KANMX4 RAD5* | (González-Prieto et al. 2013) | 3B |
| Wsgs1m2-2C | *MATa leu2-3,112 trp1-1 ura3-1 ade2-1 his3-11,15 can1-100 BAR1 pep4Δ::ADE2 sgs1Δ::KANMX4 mcm2-3A (hphNT) RAD5* | This work | 3B |
| Wsgs1d3-3C | *MATa leu2-3,112 trp1-1 ura3-1 ade2-1 his3-11,15 can1-100 BAR1 pep4Δ::ADE2 sgs1Δ::KANMX4 dpb3Δ::KANMX4 RAD5* | This work | 3B |
| Wsgs1m2d3-12A | *MATa leu2-3,112 trp1-1 ura3-1 ade2-1 his3-11,15 can1-100 BAR1 pep4Δ::ADE2 sgs1Δ::KANMX4 mcm2-3A (hphNT) dpb3Δ::KANMX4 RAD5* | This work | 3B |
| Wsgs1r5-4A | *MATa leu2-3,112 trp1-1 ura3-1 ade2-1 his3-11,15 can1-100 BAR1 sgs1Δ::KANMX4 rad5-535* | This work | 3B |
| Wsgs1m2r5-7A | *MATa leu2-3,112 trp1-1 ura3-1 ade2-1 his3-11,15 can1-100 BAR1 sgs1Δ::KANMX4 mcm2-3A (hphNT) rad5-535* | This work | 3B |
